# Vi-TIS: a transposon insertion sequencing framework for quantitative discrimination of viable and non-viable mutants

**DOI:** 10.64898/2026.05.28.728364

**Authors:** Jessica R Rooke, Karthik Pullela, Emily C A Goodall, Adam F. Cunningham, Daniela Vollmer, Waldemar Vollmer, Ian R. Henderson

## Abstract

Transposon-insertion sequencing (TIS) is a high-throughput approach that uses randomly generated transposon mutant libraries to assess bacterial gene essentiality and their contribution to fitness under defined conditions. The method integrates classical transposon mutagenesis with next-generation sequencing. However, conventional TIS workflows generally require at least one active growth step to enrich for replicating or fit mutants. The outcome of these experiments can be biased by the recovery of non-replicating cells or by the loss of slow-growing mutants. To overcome this limitation, we modified the traditional TIS workflow by incorporating propidium monoazide (PMA). PMA is membrane-impermeant and excluded from viable cells but can enter dead cells when the membrane has lost its integrity. PMA covalently crosslinks to DNA upon exposure to UV light. Such binding inhibits PCR amplification, an essential step in TIS library preparation. This property enables selective exclusion of DNA originating from dead cells and reduces the signal to noise ratio in TIS experiments. We refer to this modified approach as Viability-TIS (Vi-TIS). Applying Vi-TIS to our ultra-dense *Escherichia coli* K-12 BW25113 TIS library enhanced the accuracy of essential gene identification. We further exposed the library to subinhibitory concentrations of carbenicillin and found that Vi-TIS produced results consistent with those obtained using culture-based methodologies. In addition, Vi-TIS revealed previously unrecognized carbenicillin-susceptible mutants. Finally, we demonstrate that the mutations are linked to perturbations in peptidoglycan biosynthesis.

## Introduction

Transposon insertion sequencing (TIS) integrates classical transposon mutagenesis with next-generation sequencing (NGS) to enable genome-wide analysis of gene essentiality and fitness (Langridge et al. 2009; Cain et al. 2020; Warner et al. 2023). By quantifying the relative abundance of mutants within highly complex transposon libraries, TIS provides a powerful means of linking genotype to phenotype under defined experimental conditions (Langridge et al. 2009). This approach has been instrumental in delineating essential gene sets across diverse organisms (Langridge et al. 2009; Goodall et al. 2018; Goodall et al. 2023; Stoakes et al. 2023; A. Ghomi et al. 2024; F et al. 2024), assessing gene fitness in response to chemical agents such as antibiotics and pxidative stress (Yasir et al. 2020; Goodall et al. 2021; Roth et al. 2022; Wang et al. 2024), and probing bacterial responses to biological stresses including serum, pH, and urine (Phan et al. 2013; Gray et al. 2024; Antunes et al. 2025). TIS has been widely used to characterize bacterial physiology (Goodall et al. 2021; Winkle et al. 2021; Rooke et al. 2024; Yang et al. 2025) and has also facilitated insights into fitness determinants for *in vivo* infections (Chaudhuri et al. 2013; Karlinsey et al. 2019).

Beyond traditional implementations, several methodological innovations have expanded the utility of TIS. These include the use of modified transposons with outward-facing inducible promoters (TraDIS-Xpress; (Yasir et al. 2020)), fluorescence-activated cell sorting (FACS) to separate mutants independently of growth-based selection (TraDISort; (Paulsen et al. 2017)), and selection systems restricting detection to insertions within actively translated coding sequences (PIRT-Seq; (Goodall et al. 2025)). Despite these advances, most TIS workflows continue to rely on bacterial culturing both for library construction and for differentiating fit from less-fit mutants. These culture-dependent steps increase experimental duration and can introduce growth-related biases. For instance, dead cells may be retained during sample processing, leading to sequencing reads that confound essentiality analyses; similarly, slow-growing mutants may be erroneously classified as essential due to their apparent depletion. While approaches such as TraDISort mitigate some of these issues, FACS-based workflows are technically demanding, require specialized expertise, and often necessitate sorting millions of events to achieve adequate library coverage. Given these limitations, we sought to develop a rapid and robust method to reduce signal-to-noise ratios and improve the accuracy and efficiency of TIS experiments.

Another separate methodology is viability PCR (vPCR). This was developed as an amplification-based methodology to differentiate between live and dead cells. vPCR negates the impact of extracellular DNA, and DNA from dead bacteria, by the action of compounds such as propidium monoazide (PMA) (Nocker et al. 2006). PMA is a derivative of propidium iodide (PI), a dye commonly used to stain non-viable cells during fluorescence-activated cell sorting (FACS) or in live/dead microscopy. Both PI and PMA are excluded from intact bacterial cells, as they poorly permeate the cell envelope of viable bacteria and are further limited by active efflux (Nocker et al. 2006). However, in cells with compromised membranes, such as dead cells, both PMA and PI can enter the cell and intercalate into double-stranded DNA (Nocker et al. 2006). In contrast to PI, upon exposure to ultraviolet light (UV), PMA forms irreversible covalent bonds with the DNA. Such covalently modified DNA cannot be denatured during PCR and therefore is not amplified and therefore, only DNA from viable cells that is not bound by PMA can be amplified by PCR.

We hypothesized that incorporating PMA into the TIS amplicon-sequencing workflow would enable discrimination between live and dead bacteria in a culture-independent manner, thereby reducing signal-to-noise and increasing the sensitivity of TIS for detecting essential and conditionally essential genes. We refer to this modified approach as Viability-TIS (Vi-TIS) and the rationale is outlined in Figure 1. Here we demonstrate that PMA effectively differentiates live from non-viable mutants both in an untreated (“neat”) TIS library and under stress conditions such as antibiotic exposure. Application of PMA to the neat library improved the sensitivity with which certain mutants could be classified as essential. Moreover, incorporation of PMA during exposure to sub-inhibitory concentrations of carbenicillin accurately predicted mutant susceptibility to this antibiotic. Using this approach, we identified *sucB* mutants as highly susceptible to carbenicillin. This gene encodes an enzyme of the tricarboxylic acid (TCA) cycle responsible for generating succinyl-CoA, a precursor in several biosynthetic pathways. We show that *sucB* mutants are deficient in the peptidoglycan (PG) precursor meso-diaminopimelate (mDAP), and that supplementation of *m*DAP restores their ability to tolerate sub-inhibitory carbenicillin. These findings reveal an unappreciated functional link between *sucB*, *m*DAP biosynthesis, and β-lactam susceptibility, relationships that would likely have been overlooked using conventional TIS methods.

**Figure 1.**
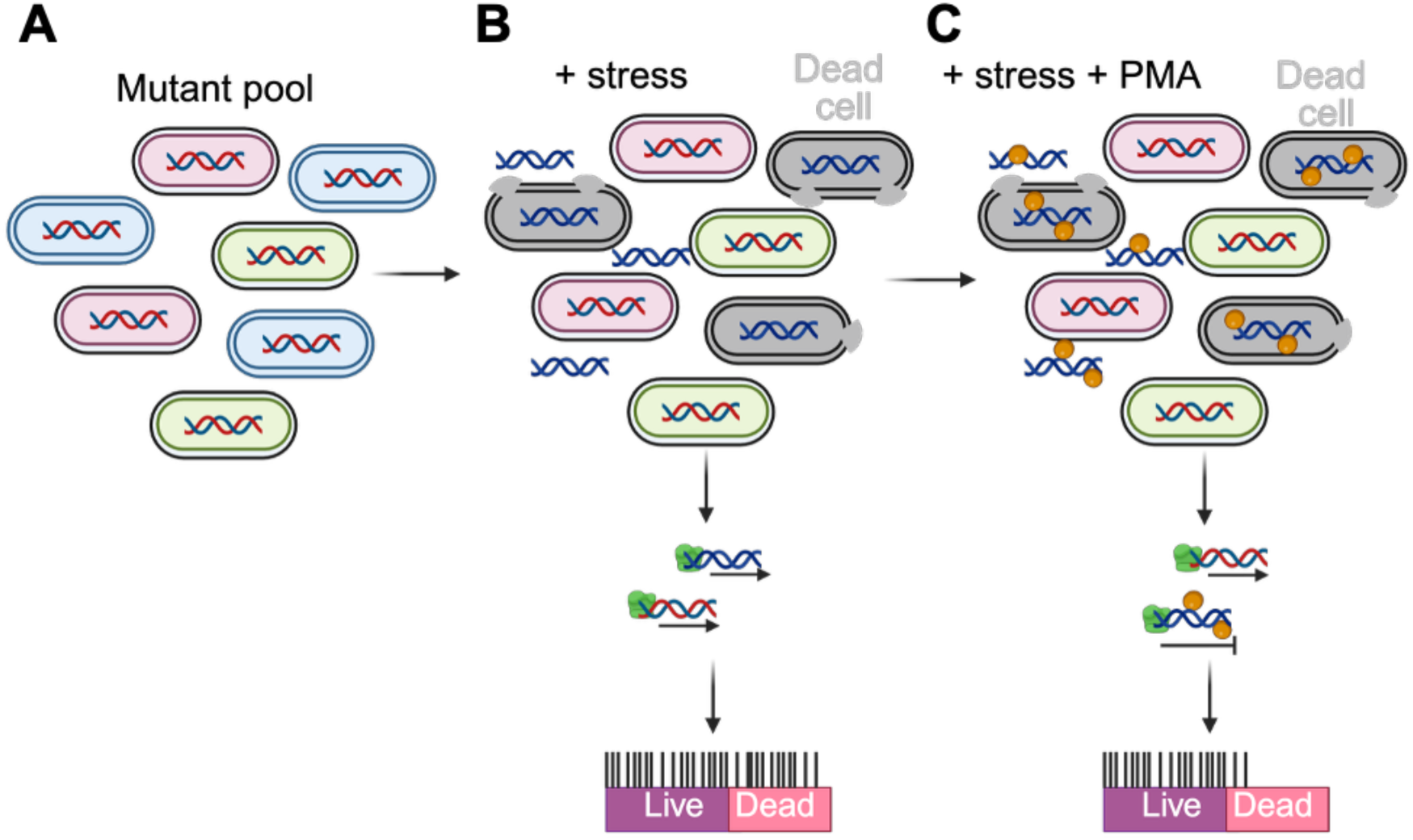
ViTIS schematic diagram. (*A*) Pool of transposon mutants. (*B*) PCR amplification of mutant DNA after exposure to stress in the absence of PMA. Both DNA from live and dead cells will be amplified resulting in identification of both live and dead transposon mutants in TIS outputs. (*C*) PCR amplification of mutant DNA in the presence of PMA. DNA from dead cells is covalently bound by PMA and PCR is inhibited. The “dead” mutants are then not identified in the TIS outputs.

## Results

### Propidium monoazide treatment for selecting viable TIS mutants

To determine whether cross-linked PMA inhibits PCR amplification from nonviable cells, we tested genomic DNA (gDNA) extracted from *E. coli* BW25113 under five conditions: (i) live cells, (ii) live cells incubated with 20 mM PMA prior to UV exposure, (iii) dead cells, (iv) dead cells incubated with PMA, and (v) dead cells incubated with PMA followed by UV. Nonviable cells were generated by incubating cultures in 80% (v/v) ethanol for 5 min, followed by PBS washes. Using primers targeting *bamE*, which encodes a conserved component of the outer membrane protein biogenesis machinery (insert appropriate reference), we observed a PCR product for all conditions except dead cells treated with PMA and exposed to UV (Fig. S1A). These results demonstrate that PMA crosslinking selectively suppresses amplification from dead, but not live, *E. coli*. To exclude UV-dependent viability artifacts, we quantified colony-forming units (CFU) with and without UV exposure and detected no significant differences (Fig. S1B).

Having established that PMA treatment, followed by UV exposure, effectively prevents PCR amplification of DNA from non-viable cells, we next assessed the impact of PMA treatment on a TIS library in *E. coli* BW25113. This library, constructed using a mini-Tn5 transposon carrying a chloramphenicol resistance cassette, has been described previously (Goodall et al. 2018). Previous sequencing revealed the library contained 901,383 Unique Insertion Points (UIPs) but that 199,557 UIPs (22%) were represented by one read (Goodall et al. 2018), raising the possibility that these low-abundance insertions originated from dead cells. To test this hypothesis, an aliquot of the library was thawed on ice, treated with 20 mM PMA, and exposed to UV. Genomic DNA was then extracted, transposon-genome junctions were PCR-enriched, and sequencing was performed, as described previously (Goodall et al. 2018). Raw reads were processed to remove tags, and transposon insertion sites were identified using BioTraDIS (Barquist et al. 2016). To assess reproducibility, we compared gene-level insertion counts between replicates for each condition, which yielded high correlation coefficients for both conditions (0.99 and 0.96; Fig. S2). Consequently, replicate datasets were combined, resulting in >4.8 million mapped reads for the untreated library (NTL) and >4.4 million mapped reads for the PMA-treated NTL (Table S1). Chromosomal insertion profiles for both conditions showed the expected high-density, genome-wide coverage indicating the distribution of insertions was not affected by PMA treatment (Fig. 2A). We next quantified the number of UIPs with 0-5 reads. Despite the similar sequencing depths, PMA treatment of the library resulted in a marked reduction in the number of identified UIPs from 685,977 to 429,901 (Fig. 2B-C; Table S1). We observed that PMA treatment led to a substantial increase (>5%) in genomic positions lacking detectable insertions (Fig. 2D) and a corresponding decrease in positions supported by only one or two reads (Fig. 2E). Together, these findings demonstrate that PMA treatment effectively reduces the prevalence of low-read insertion sites, supporting the hypothesis that there are a number of dead mutants within the original library.

**Figure 2.**
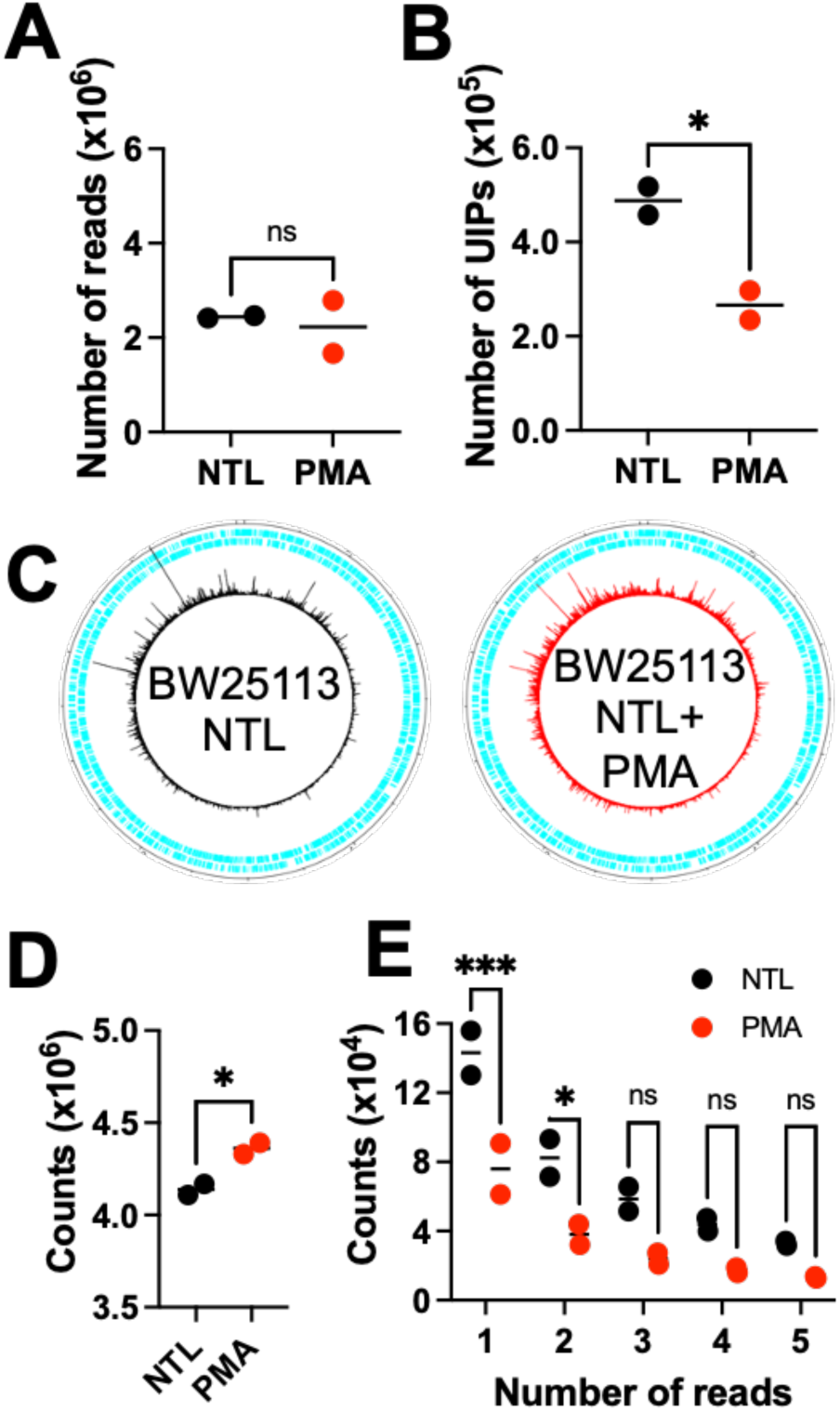
Analysis of *E. coli* BW25113 TIS library treated with and without PMA. (*A*) Circular plots showing transposon distribution across the genome in the neat library (NTL) (left) and PMA (right) samples. (*B*) Number of mapped reads for each replicate for NTL and NTL treated with PMA. (*C*) Number of unique insertion points (UIPs) in NTL and PMA samples. Number of positions in the genome that have 0 (*D*) and 1-5 (*E*) transposon insertions in each condition. Statistical significance (p<0.05) was determined by unpaired *t*-tests (panels *A*, *B* and *D*) or a 2-way ANOVA with Sidaks correction for multiple comparisons (panel *E*). * p<0.05 and *** p<0.001.

### PMA treatment alters the number of genes called as “essential”

Having established that PMA treatment effectively reduces the contribution of reads derived from non-viable cells to TIS datasets, we next sought to determine how this reduction influences our ability to identify essential genes. To investigate this the frequency of insertions along each coding sequence was first normalised by gene length and genome size to generate an insertion index score for each condition. Both NTL and PMA-treated samples formed a bimodal distribution (Fig. S3A), but there was an overall shift in the PMA-treated sample, where the median insertion index score decreased from 0.146 (NTL) to 0.0859 (Fig. S3B), and this was consistent across all the genes on the genome (Fig. S3C). Taken together, these data suggest that PMA treatment reduces the recovery of low-abundance UIPs across the genome, thereby sharpening the statistical threshold for distinguishing essential from non-essential genes and increasing confidence in the resulting data. To ensure that PMA did not overly bias our data, we compared the similarity of data by plotting the number of insertions per gene of the NTL and PMA-treated samples against each other. We observed a good correlation with the PMA-treated sample and the untreated NTL (r > 0.95) indicating that even though there was a decrease in the number of UIPs, this did not bias the analysis (Fig. S3D). We next assayed the number of essential genes in both conditions. Using the BioTradis essentiality.R script as well as manual confirmation using the artemis genome browser software. We determined that 372 genes were essential in the NTL, similar to previous publications (Goodall et al. 2018). We also identified an additional 28 genes that were called as essential only in the PMA-treated condition (Fig. 3A and Table. S2). We first confirmed that these extra essential genes were not mutants that were known to create permeability defects in the bacterial envelope, or known to affect efflux, such as *surA* (Justice et al. 2005)*, yhcB* (Goodall et al. 2021)*, tolC* and *acrAB* (Blair et al. 2014) (Fig. 3B), suggesting that the concentration of PMA used in the experiment (20 mM) does not introduce notable biases into the data.

**Figure 3.**
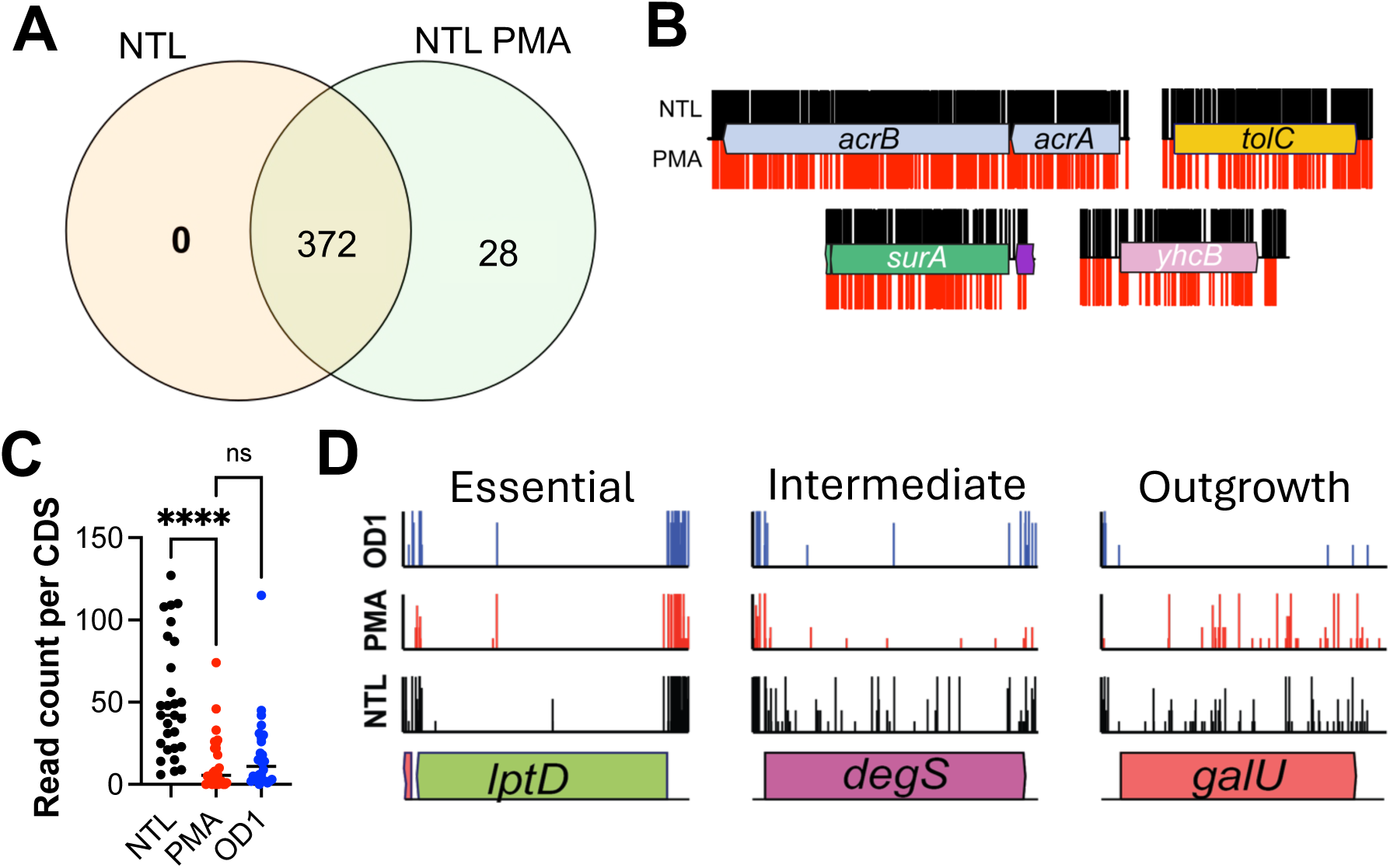
The effect of PMA treatment on essential gene analyses. (*A*) Venn diagram comparing the number of essential genes identified in NTL and PMA. (*B*) Insertion plots of mutants with known cell envelope defects in both NTL (top, black) and PMA (bottom, red). (*C*) Read counts for the 28 genes found to be uniquely essential in the PMA-treated library compared to the readcounts for the same genes in the NTL and LB outgrowth samples. Statistical significance (p<0.05) was determined using One-way ANOVA with Tukeys correction for multiple comparisons. **** p<0.0001. (*D*) Examples insertion plots of always essential, intermediate and outgrowth essential genes.

Prior work indicated that transposon mutants in certain essential genes can persist through additional rounds of cell division relative to others, depending on factors such as protein abundance or function (Gallagher et al. 2020). Based on these observations, we hypothesized that the additional 28 genes identified as essential in the PMA-treated sample represent non-viable mutants within the library; specifically, mutants capable of undergoing several rounds of cell division before loss of viability. To evaluate this possibility, we compared our set of 28 genes to genes classified as essential following outgrowth of the library to an OD_600nm_ of 1 in LB medium. The insertion index values for the 28 PMA-specific essential genes were comparable to those observed when the library was cultured to an OD_600nm_ of 1 (Fig. 3C). These findings suggest that the 28 genes are indeed lost from the population and that this loss likely occurs between library preparation, storage, and subsequent outgrowth. Therefore, we classified these loci as “intermediate essential” genes, reflecting the current uncertainty regarding the specific stage at which their corresponding mutants lose viability. Our data support a core set of 372 essential genes, an additional 28 intermediate essential genes, and a further 12 genes that become non-viable upon growth in liquid culture to an OD_600nm_ of 1 (Fig. S4), with representative examples shown in Figure 3D. Collectively, these results indicate that loss of mutant viability is more nuanced than typically appreciated and that conventional TIS approaches, whether using a neat or outgrown library, would not satisfactorily resolve this complexity without the incorporation of PMA treatment.

### Effect of carbenicillin on *E. coli* BW25113 mutant library

We have established the PMA can be used in conjunction with TIS to improve the signal to noise ratio. We hypothesised that the inclusion of PMA into the TIS workflow would improve the identification of genes required for growth under conditions of stress. To test this hypothesis, we exposed our transposon library to a sub inhibitory concentration of carbenicillin with and without PMA. First, using an endpoint assay, we established that the minimum inhibitory concentration (MIC) of carbenicillin for *E. coli* BW25113 was 16 μg/ml (Fig. S5A). We selected a sub-inhibitory concentration corresponding to one-eighth of the MIC (2 µg/ml) and confirmed that this dose did not alter the growth dynamics of the wild-type bacterium (Fig. S5B) or reduce viable colony-forming units (CFU) after 3 h of incubation (Fig. S5C). To identify mutants exhibiting heightened susceptibility to β-lactam treatment, we exposed approximately 10⁷ CFU/ml of the transposon mutant library to 2 µg/ml carbenicillin for 3 h, in duplicate. After 3 h of exposure, one set of samples were directly processed for gDNA extraction, one was treated with PMA and a third was sub-sampled and re-inoculated into LB medium lacking carbenicillin and grown until an OD_600nm_ of 1 before gDNA extraction. After gDNA was harvested, samples were prepared for sequencing according to our bespoke protocol for TIS, as outline previously (Goodall et al. 2018). For each sample, we mapped between 5.6-7.3 million reads, which corresponded to 640,000-770,000 UIPs (Table S3). To assess the reproducibility of each experiment, we compared gene-level insertion index scores between replicates for every sample. This analysis revealed strong concordance for each condition, with correlation coefficients ranging from 0.90 to 0.99 (Fig. S5D–F). Having established reproducibility, we merged the replicate datasets and using the Bio-TraDIS tradis_comparisons.R script we quantified gene fitness under each condition by comparing changes in read counts per gene to the LB outgrowth dataset. Genes exhibiting significantly reduced fitness were defined by a q-value < 0.05 and a log₂ fold change (logFC) less than -2. Applying these thresholds, we identified 8 genes with decreased fitness after 3 h of exposure to sub-inhibitory carbenicillin, 18 genes after 3 h of exposure combined with PMA treatment, and 24 genes following the regrowth condition (Fig. 4A–C and Tables S4, S5 and S6).

**Figure 4.**
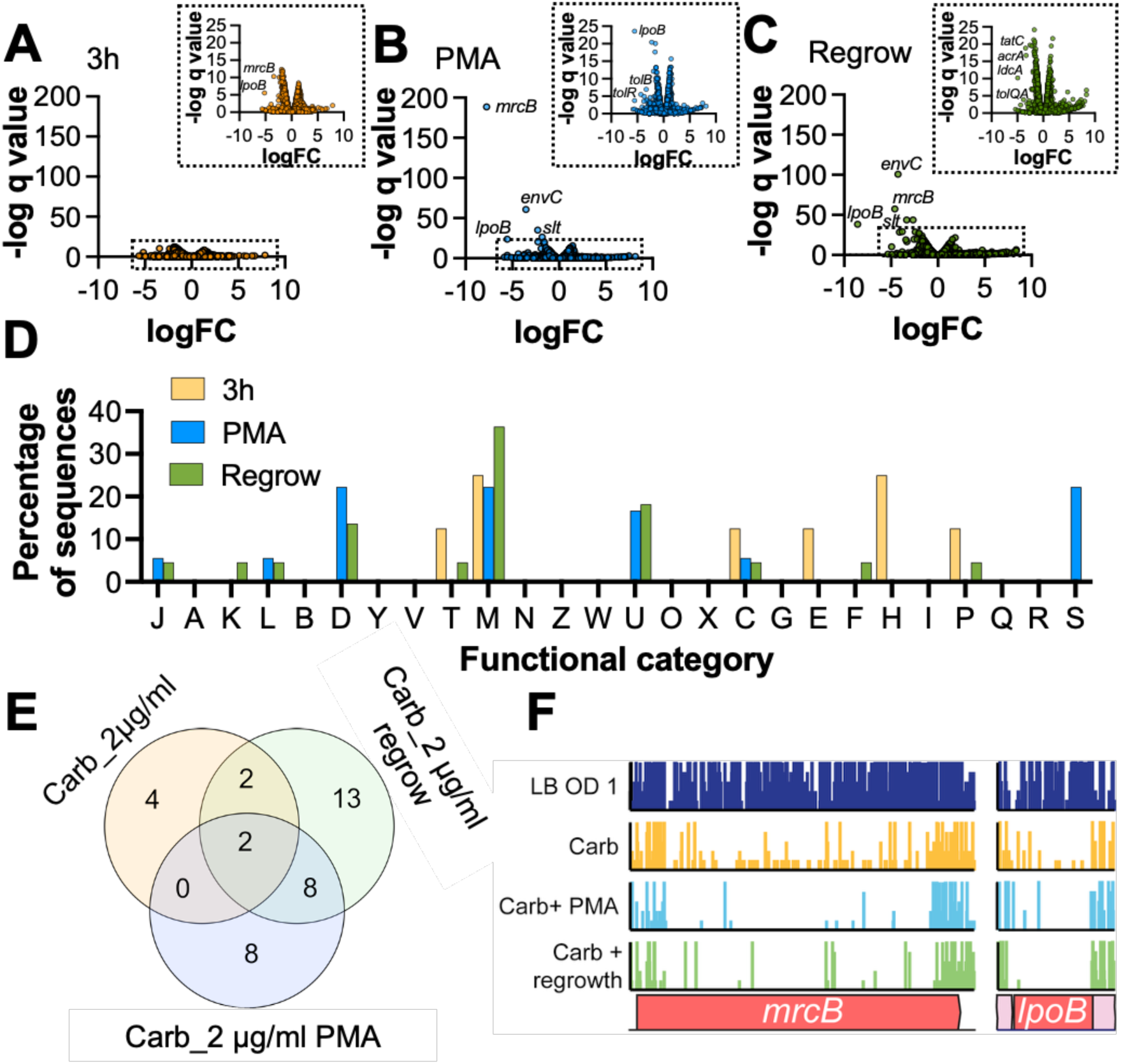
Carbenicillin exposure identifies known targets for β-lactam antibiotics. Fitness of TIS mutants after 3 h of exposure to 2 μg/ml carbenicillin (*A*), 3 h of exposure to 2 μg/ml carbenicillin with PMA treatment (*B*) and 3 h of exposure to 2 μg/ml carbenicillin with subsequent re-inoculation into antibiotic free LB medium grown to an OD_600nm_ of 1 (*C*). (*D*) COG analysis of identified susceptible mutants (q value <0.05 and logFC < -2). Definitions for COG letters are defined in Table. S7. (*E*) Venn diagram comparing unique and overlapping mutants identified in each screening condition. (*F*) Insertion plots capped to five reads of the *mrcB* and *lpoB* loci for each condition.

Having established that each experiment revealed different sets of genes associated with decreased fitness in the presence of carbenicillin, we analysed the functional category of the identified genes using Clusters of Orthologous Genes (COG) analysis (Tatusov et al. 2000). We identified that for each condition over 20% of genes were assigned the COG category M (cell wall/membrane/envelope biogenesis) (Fig. 4D). In addition, both PMA-treated and outgrowth samples identified genes with functional categories D (cell cycle control, cell division, chromosome partitioning) and U (intracellular trafficking, secretion and vesicular transport) (Fig. 4D). These data indicate that the identified mutants possess functional annotations consistent with known β-lactam responses, although carbenicillin exposure alone yielded the smallest number of genes within the expected functional categories (Fig. 4D).

To better understand the outcome of these screening experiments we assessed which genes were unique and shared for each condition. We observed that PMA-treated and the outgrowth samples had the largest number of overlapping genes (8), and each had 8 and 13 uniquely identified genes, respectively (Fig. 4E). Transposon mutants in the *lpoB* and *mrcB* genes were found to be less fit in all three conditions when compared to untreated controls (Fig. 4E-F). Both genes are known to be susceptible to β-lactam antibiotics, with *mrcB* encoding penicillin binding protein 1B (PBP1B) and LpoB its outer membrane-anchored activator (Typas et al. 2010). In addition, we identified within the 8 genes shared between PMA-treated and outgrowth samples, genes known to be more susceptible to β-lactam antibiotics. These included mutants **Figure 4** lacking *envC*, *slt*, or *tol-pal* genes (Zahir et al. 2020; Kimura et al. 2025) (Fig. S6). Our findings demonstrate that the PMA screen effectively identified genes known to be associated with β-lactam tolerance.

### PMA treatment successfully identifies carbenicillin susceptible mutants

Having established that the PMA screen can identify genes associated with carbenicillin susceptibility, we sought to assess the accuracy of each experimental condition in identifying such mutants. To do this, we evaluated the corresponding kanamycin resistant mutants from the *E. coli* BW25113 Keio single-gene deletion library (Baba et al. 2006) in a competitive growth assays to determine their relative fitness in the presence of carbenicillin. Of the mutants identified in our screen, 30 of 37 had a matched Keio mutant strain. Each mutant was mixed with the parent strain in a 1:1 ratio before inoculation into LB broth or L-broth containing 2 µg/ml carbenicillin, followed by incubation for 3 h at 37°C. Viable CFU were then enumerated by plating serial dilutions on LB agar supplemented with and without kanamycin, to distinguish between the number of viable wild-type and mutant bacteria. A competitive index was calculated for each mutant. While 4 of the 30 mutants had a defect in growth compared to the wildtype in LB broth alone, 22 of the 30 mutants were significantly outcompeted by the parent strain when treated with carbenicillin (Fig. 5A). We next compared TIS-derived fitness scores with fitness outcomes from the competitive assays. Carbenicillin exposure alone showed the lowest predictive accuracy for identifying susceptible mutants, failing to detect 82% of the 22 susceptible mutants (Fig. 5B and 5E). In contrast, PMA-treatment and the outgrowth conditions exhibited stronger agreement (*r* = 0.57 and *r*= 0.62, respectively; Fig. 5C, D and E), producing the highest validation success rates (90% and 86%, respectively) and very low false-positive rates (Fig. 5E).

**Figure 5.**
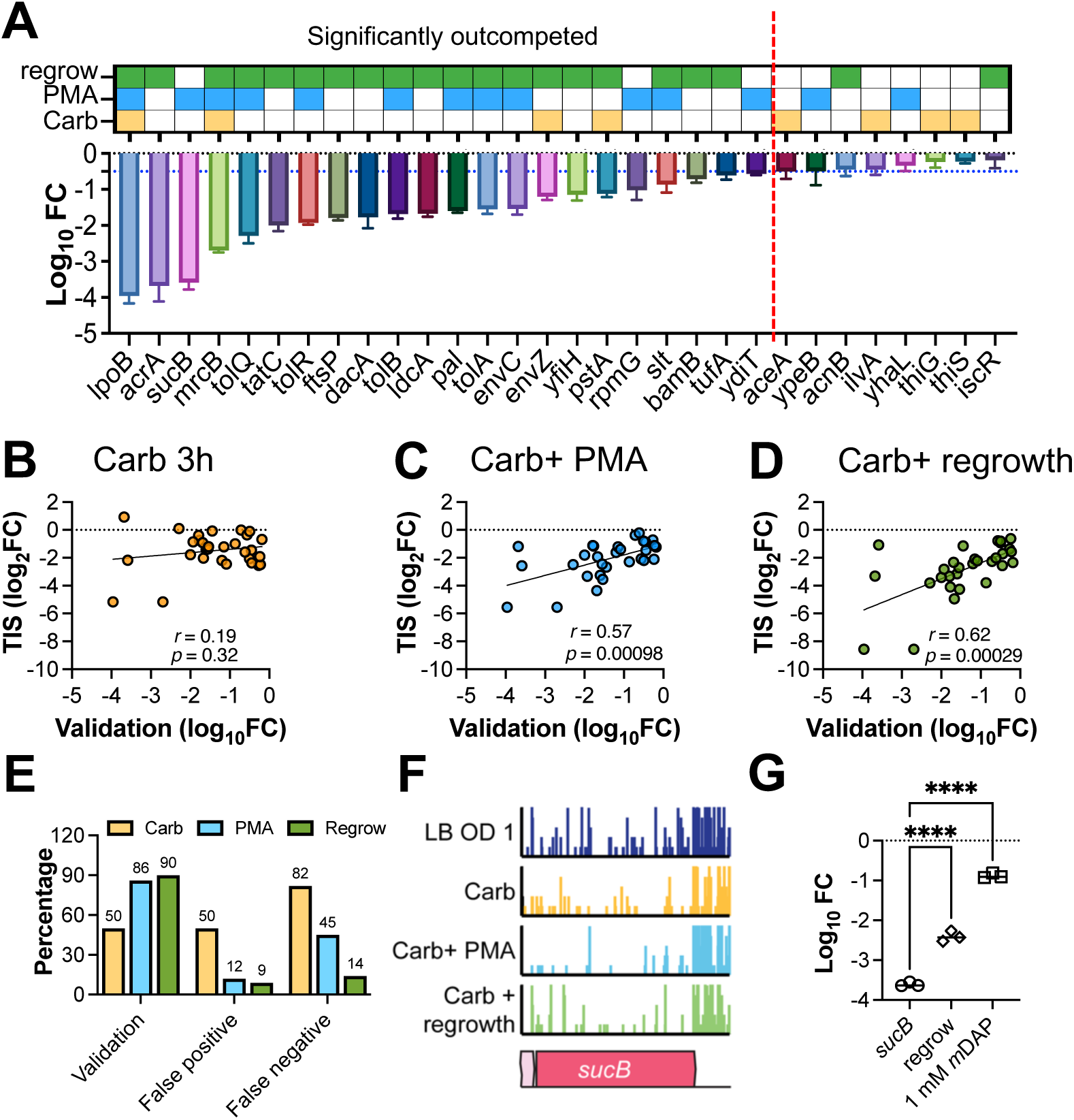
TIS validations and role of *sucB* in carbenicillin tolerance. (*A*) Bar chart showing the Log_10_ fold change (FC) in cfu between competitive growth assays (1:1 of WT and mutant) in LB and LB with 2 μg/ml of carbenicillin. Significance (*p* < 0.05) was determined by One sample *t*-test and a mutant considered significantly outcompented if the *p* value was <0.05 and the log_10_ FC <-0.5. (*B*-*D*) Scatter plots comparing Log_2_FC of mutants predicted in TIS vs Log_10_FC of Keio mutants during validation competeitve growth assays with (B) showing fitness prediction from carbenicillin exposure (carb 3 h), (*C*) carbenicillin exposure with PMA (Carb + PMA) and (*D*) carbenicillin exposure and regrowth (Carb + regrowth). (*E*) Graph showing validation, false positive and false negative percentages for each condition. (*F*) Insertion plot (capped at 5 reads) of the *sucB* loci in all conditions. (*G*) Fitness of *sucB* mutants after 3 h of carbenicillin exposure (*sucB*), after subsequent regrowth in antibiotic free LB medium (regrow) or with the addition of 1 mM *meso*-diaminopimelate (1 mM *m*DAP). Statistical significance (p<0.05) was determined using a one-way ANOVA with Dunnetts correction for multiple comparisons **** p<0.0001.

Together, these results demonstrate that both PMA treatment and the outgrowth conditions provide accurate and reliable predictors of mutant susceptibility to carbenicillin.

### PMA treatment identifies new genes involved in β-lactam tolerance

During our validations, we observed that similar to *lpoB* and *mrcB, sucB* mutants were highly outcompeted during exposure to carbenicillin (Fig. 5A), but *sucB* was only identified in the PMA condition (Fig. 5A and F). We hypothesised that upon re-inoculation into LB (regrowth), *sucB* mutants may recover from the initial carbenicillin stress. To test this, we first inoculated *sucB* mutants and BW25113 WT 1:1 into LB medium with 2 μg/ml carbenicillin for 3 h. We then plated viable CFUs onto LB agar and also re-inoculated a subset of the culture into fresh LB without antibiotics. We then incubated this culture until an OD_600nm_ of approximately 1 and again plated viable CFUs. We identified that prior to re-inoculation, the *sucB* mutants had 3.5 log_10_ reduction in CFU in carbenicillin compared to LB controls. On re-growth, this had significantly reduced to a 2.5 log_10_ difference (Fig. 5G). These data demonstrate that re-inoculation into LB medium significantly increased *sucB* mutant viability after initial antibiotic exposure and would provide an explanation as to why *sucB* was not identified in the regrowth TIS screen.

*sucB* encodes a component of the 2-oxoglutarate dehydrogenase enzyme complex that is part of the tricarboxylic acid (TCA) cycle (Smith and Neidhardt 1983; Cronan and Laporte 2005; Karp et al. 2025) (Fig. S6). SucB, in conjunction with SucA, converts 2-oxoglutarate into succinyl-CoA, which is then converted into succinate via SucCD (Cronan and Laporte 2005; Karp et al. 2025) (Fig. S7). In our data, only *sucB* was significantly susceptible to carbenicillin, suggesting that production of Succinyl-CoA is important for antibiotic tolerance. Succinyl-CoA feeds into multiple metabolic pathways; one being the TCA cycle and the other being the L-lysine biosynthetic pathway, where it is used by DapD to convert *N*-succinyl-2-amino-6-ketopimelate into *N*-succinyl-L,L-2-6-diaminopimelate (Karp et al. 2025) (Fig. S7). The L-lysine biosynthetic pathway generates multiple products, including *meso*-diaminopimelate (*m*DAP) (Davis 1952; Karp et al. 2025). *m*DAP is the third peptide in the stem peptide chain of PG in *E. coli* and is the site for PG crosslinks (Garde et al. 2021). We hypothesised that *sucB* mutants would have reduced cellular levels of *m*DAP and this in turn would result in fewer PG crosslinks and an increased susceptibility to β-lactam antibiotics. Therefore, to test this hypothesis, we added *m*DAP to the growth medium and tested the competitive index of *sucB* mutant vs BW25113 WT in the presence of carbenicillin. Addition of *m*DAP completely restored *sucB* mutant tolerance to carbenicillin (Fig. 5G), suggesting the susceptible phenotype of this mutant was due to reduced cellular levels of *m*DAP.

## Discussion

Distinguishing between live and dead bacterial cells is essential to ensure that measurements and analyses reflect only biologically active populations, enabling accurate interpretation of function, genetic fitness, and clinical interpretation. As such, multiple methodologies have been developed to try to define and measure these proportions in bacterial populations (Boulos et al. 1999; Davey 2011; Cangelosi and Meschke 2014). In so doing, we often make assumptions on the definitions of what is considered “alive” and what is considered “dead” and what are the parameters used within a study to measures these phenomena. One such methodology is vPCR, which has been used to understand viability of bacterial populations in the environment (Delgado-Viscogliosi et al. 2009; Letant et al. 2011; Nisar et al. 2023) and to ascertain viability of pathogens (Vojtech et al. 2023; Veugen et al. 2024). PMA selectively binds to extracellular DNA or DNA from dead cells that have compromised cell membranes (Nocker et al. 2006), although the precise mechanism of action of PMA is not yet completely understood. In addition to preventing denaturation of DNA when covalently bound, experiments suggest that the poor solubility of the PMA bound DNA results in precipitation of the genomic DNA leading to further exclusion of gDNA from membrane compromised dead bacteria during DNA purification step (Nocker et al. 2006; Tekgul and Adiguzel 2025).

TIS data analysis relies on the relative abundance of transposon mutants within a pooled library to predict essentiality and/or fitness of mutants under conditions of interest (Cain et al. 2020; Warner et al. 2023). There is an overall assumption that the presence of a transposon mutant in a population represents that mutation as being “viable” and the cells containing it are “alive”. Whereas the consistent absence or reduction of a transposon mutant in a population assumes that cells containing that mutation are either “dead” or outcompeted by the rest of the population. The key to a successful TIS screen is the ability to differentiate between the fit/living mutants versus the dead/unfit ones. But given TIS methodologies rely on amplification of transposon-genome junctions via PCR, even DNA from dead cells could still contribute to data, thereby obscuring or limiting the screen in its statistical power. Growth enrichment is one such way of increasing the chances that the abundance of transposon insertions from living cells massively outnumber those from dead cells. However, one of the primary concerns while studying TIS libraries under stress using a culture-dependent enrichment is the possible recovery of susceptible mutants under non-selective growth conditions, which could result in overestimating the fitness or completely fail to identify susceptible mutants. We directly observed this phenomenon in our study where *sucB* mutants were able to partially recover after carbenicillin exposure when re-inoculated into antibiotic free medium. Secondly, while calling essential genes, mutants with reduced growth rate could be under-represented in the population resulting in lowered relative abundance leading to false positives. Thirdly, mutants that are more fit under ‘perturbing’ growth conditions would be less obvious in the TIS study upon regrowth of the mutant library under non-perturbing conditions. Finally, the current methodologies fail to give nuances of bacteriostatic and bactericidal effects as both these will only be labelled as less fit.

In this study, we utilised the addition of PMA to traditional TIS methodology to differentiate between live and dead mutants in both the NTL and the library exposed to sub-inhibitory concentrations of carbenicillin. By using PMA in this way, we were able to establish that PMA does not bias the library by over-targeting mutants with disrupted cell membrane and the addition of PMA provided some resolution of mutants that lose viability somewhere between making the library, storage and outgrowth. Our carbenicillin screen identified genes known to be important for β-lactam tolerance, including *mrcB, lpoB, slt, envC, tol-pal*, *dacA*, *ldcA* and *ftsP* (Phan et al. 2020; Zahir et al. 2020; Kimura et al. 2025), with an overall validation rate of approximately 73%. Usage of PMA after antibiotic stress resulted in high (86%) identification of true susceptible mutants and showed comparable results to the growth enriched sample, demonstrating that PMA could be used in similar screens and would provide similar outputs as growth enrichment but without the need for an additional growth step. In addition, PMA usage identified a mutant that recovered under the stress-free regrowth conditions (*sucB*). Previous studies have identified that *sucB* mutants are importance for persister formation in the presence of multiple antibiotics (Ma et al. 2010), but a mechanism for this phenotype was not previously elucidated.

The metabolic product produced by *sucB*, as part of a larger enzyme complex, is succinyl-CoA. Previous studies have linked the over-abundance of succinyl-CoA with an overproduction of L-lysine in *Corynebacterium glutamicum* (Kind et al. 2013). L-lysine is one of the products synthesised in the L-lysine biosynthesis pathway that also generates *m*DAP and that succinyl-CoA directly feeds into this pathway as a co-factor for the reaction mediated by DapD (Fig. S6). A reduction in succinyl-CoA in *sucB* deficient cells would likely lead to reduced *m*DAP levels and therefore a PG cell wall with fewer cross-links. In this study, we identified that *sucB* mutants are deficient in *m*DAP and this is likely the mechanism behind β-lactam susceptibility as opposed to the role of *sucB* during energy production and central carbon metabolism.

We envision that Vi-TIS could be an alternative method to remove or reduce the need for a growth enrichment step during TIS screening. This could save time and can significantly reduce the handling and experimentation time while providing more insight into the susceptible mutants under the study conditions. Vi-TIS could be a cheaper and user-friendly alternative to TraDIS-Sort as a method to differentiate between live and dead mutants. Vi-TIS could also have utilisation in scenarios that are known to produce extracellular DNA, such as in biofilm conditions (Montanaro et al. 2011). In this study we show that Vi-TIS can accurately identify highly susceptible mutants even when using low concentrations of antibiotic (1/8^th^ MIC), which could be useful if a compound is expensive or hard to procure or produce. In summary, Vi-TIS is a novel variation on traditional TIS that can differentiate between live and dead mutants in a culture-independent manner, and this study provides the first proof of principle demonstration of its uses and potential.

## Methods

### Standard culture conditions

Bacteria were grown in either liquid or solid (1.5% agar) Luria-Bertani (LB) medium unless stated otherwise. Media was supplemented with kanamycin (50 μg/ml) or carbenicillin (various concentrations) as required. Unless stated otherwise, bacteria were incubated at 37°C with shaking. Strains used in this study are outlined in Table S8.

### Molecular techniques

For PCR amplification of DNA, MyTaq redmix (Bioline) was used following manufacturers guidelines. DNA was analysed using 1% agarose gels and SYBR safe (Thermo) DNA stain. Gels were visualised using a BioRad Gel Doc.

### Minimum inhibitory concentrations (MIC)

Carbenicillin was added to Mueller-Hinton broth at concentrations ranging from 512-1 μg/ml and no antibiotic controls in a 96 well plate. Overnight cultures of *E. coli* BW25113 were washed and added to wells on the 96 well plate at a density of 5x10^5^ cfu per well, verified by plating serial dilutions on LB agar. 96 well plates were covered with a gas permeable membrane and incubated at 37°C for 16 h. After incubation, OD6_00nm_ was measured using a Polarstar plate reader and MIC determined as the concentration of antibiotic that did not show any growth by eye.

### PMA exposure

PMA was sourced from Gene Target Solutions Pty Ltd. Approximatley, 10^9^ cfu of a bacterial sample was treated with 20 mM of PMA and subsequently exposed to UV light for 3 min following the manufacturers protocols. Following UV exposure, gDNA was extracted using the RTP Stratec DNA genome extraction kit. UV exposure and spot plates: Overnight cultures were normalised to 10^9^ cfu/ml and exposed to UV light for 3 min. Following exposure, cells were serially diluted and spot plated onto LB agar to enumerate viable cfu. For the TIS library screens, aliquots of the *E. coli* BW25113 TIS library were thawed on ice and inoculated into either LB or LB supplemented with 2 μg/ml of carbenicillin in duplicate to a starting OD_600nm_ of 0.02. This density allowed each of the approx. 1 million unique mutants in this library to be sampled approximately 1000x. The cultures were incubated at 37°C with shaking until an OD_600nm_ of 1. Bacterial cells were isolated by centrifugation and processed for gDNA extraction using the RTP Stratec DNA genome extraction kit.

### TIS sequencing

gDNA was fragmented to approximately 200 bp using a Covaris Bioruptor and transposon-genome junctions were amplified by PCR. The amplified products were prepared for sequencing on Illumina platforms using the NEB ultra 1 kit (New England Biolabs) and were quantified via qPCR using a Kapa Library Quantification kit. Sequencing reads were generated using an Illumina MiSeq and data deposited onto the European Nucleotide Archive (accession number PRJEB113234).

### TIS data analysis

Sequencing reads were demultiplexed prior to identifying reads containing the tn tag (GTCTCATTTTCGCCAAAGATGTGTATAAGAGACAG) using the BioTradis bacteria_tradis script (Barquist et al. 2016). Reads containing the tn tag were mapped to the *E. coli* BW25113 reference genome (CP009273) using Smalt with default parameters outlined in the BioTradis pipeline (Barquist et al. 2016). Transposon insertion sites were determined using the BioTradis tradis_gene_insert_sites script with a 10% trim applied to the 3’ end of reads that can often tolerate insertions even in essential genes. Essential genes were determined using the tradis_essentiality.R script that was modified to predict essential genes with a 12-fold likelihood of belonging to the essential mode of the bimodal distribution as described previously (Goodall et al. 2018). Mutant fitness was determined using the tradis_comparison.R script and the output was filtered to exclude genes that had fewer than 3.2 counts per million (CPM) reads.

### Cluster of orthologous genes (COG) analysis

The protein fasta sequence for susceptible mutants identified under each condition were classified in Cluster of orthologous genes (COGs) using COGclassifier. The data were transformed into the percentage of genes in each category and represented in a bar graph.

### Competitive index growth experiments

Overnight cultures of *E. coli* BW25113 and each mutant were normalised and added in a 1:1 ratio into LB medium with and without 2 μg/ml of carbenicillin. Cultures were incubated for 3 h at 37°C and viable bacteria were enumerated by plates serial dilations of cultures on LB agar plates. Competitive index (CI) was calculated as the number of mutant cfu/ BW25113 WT cfu in each condition. Each WT-mutant pair was repeated in biological triplicate. Significance was determined using a One-sample *t*-test determining if a mutant had a CI that was significantly (p<0.05) less than one. Log_10_ fold change (log_10_FC) was calculated by determining the difference in cfu for each mutant in LB vs LB supplemented with 2 μg/ml of carbenicillin.

### *sucB* mutant growth assays

To determine whether the *sucB* mutant recover in the absence of antibiotic stress, CI experiments were set up as described above in biological triplicate. After the 3 h incubation, viable bacteria were enumerated, and a subset of cells were centrifuged at 5,000 *x*g and washed in fresh LB broth. Fresh LB was inoculated to a starting OD_600nm_ of 0.02 of carbenicillin exposed bacteria and this was incubated at 37°C until an OD_600nm_ of approximately 1. Here, the viable bacteria were enumerated as previously described. To determine if *meso-*diaminopimelate (*m*DAP) could restore *sucB* tolerance to carbenicillin, CI experiments were set up as previously described and *m*DAP was added to a final concentration of 1 mM. After 3 h incubation at 37°C, viable bacteria were enumerated as previously described.

### Statistical analyses

Statistical significance (p<0.05) was determined using GraphPad Prism v10. For comparisons between two independent groups, significance was determined using either a paired or unpaired *t*-test. For comparisons with more than two groups or variables, we used either One-way or two-way ANOVAs with corrections for multiple comparisons.

## Acknowledgements

The authors acknowledge helpful discussions from the Henderson and Vollmer laboratories. IRH and JLR were funded by the Australian research council (DP230102796).

## Author contributions

JLR, KP and IRH conceived the project and wrote the manuscript. KP and JR acquired and analysed all of the data. ECAG edited the manuscript, assisted in data analysis and TIS expertise. AFC edited the manuscript. DV assisted in data acquisition and technical expertise. WV edited the manuscript and provided cell wall expertise. IH obtained funding for the project.

